# Cortical representation of pain by stable dedicated neurons and dynamic ensembles

**DOI:** 10.1101/2020.11.02.364778

**Authors:** Mario A. Acuña, Fernando Kasanetz, Paolo De Luna, Thomas Nevian

## Abstract

The perception of pain arises from distributed brain activity triggered by noxious stimuli. However, which patterns of activity make nociception distinct from other salient sensory experiences is still unknown. Using *in vivo* chronic two-photon calcium imaging in slightly anaesthetized mice, we identified a nociception-specific representation in the anterior cingulate cortex (ACC), that is attained by a core of neurons that code for a generalized concept of the pain experience. The overall ensemble activity allowed for an efficient discrimination of the sensory space, despite a drift in single-neuron sensory tuning over time. Following sciatic nerve lesion, the representation of nociceptive stimuli was impaired as a consequence of innocuous stimuli expanded into the nociception-specific ensemble, leading to a dysfunctional discrimination of sensory events in the ACC. Thus, the hallmark of chronic pain at the cortical neuronal network level is an impairment of pattern separation and classification identifying a circuit mechanism for altered pain processing in the brain.

## Main text

The perception of pain is a complex phenomenon that requires activity in distributed regions of the brain. However, it is still largely unclear how a noxious stimulus is processed in cortical brain regions and which patterns of activity make the pain distinct from other salient sensory events. Functional brain imaging in humans identified several brain areas activated by nociceptive stimuli, which together represent the different aspects of the pain experience [1]. Nevertheless, this activity in the so called “pain matrix” is not specific for pain. These brain regions are involved in multiple and diverse processes other than pain and similar responses in the pain matrix can be elicited by salient but non-painful sensory stimuli [2, 3]. Despite this, using functional magnetic resonance imaging, has been described that physical pain is reliably encoded by apparently specific patterns of activities across and within regions of the pain matrix [4]. Such nociception-selective responses could be achieved by dedicated local neurons embedded in a broader multifaceted neuronal network serving also other functions. Nevertheless, how neuronal ensembles inside cortical areas of the pain matrix codify nociception, as well as their degree of selectivity, is largely unknown but essential for an objective understanding of the pain experience.

The anterior cingulate cortex (ACC) is a central component of the brain response to pain and it is believed to participate in the affective and emotional connotation of nociceptive representation. The human ACC is not only reliable activated by noxious stimuli, but also the magnitude of the nociceptive response correlates with stimulus intensity and unpleasantness [5–7]. Furthermore, patients with lesions in the ACC report altered emotional pain perception[8]. In rodents, manipulations that reduce or eliminate ACC activity cause impaired responses to noxious stimuli [9–12]. However, the neuronal mechanisms by which ACC neurons encode the aversive quality of sensory stimuli and distinguish them from other salient or neutral events have not yet been characterized. *In vivo* acute and chronic electrophysiological recordings have identified ACC neuronal responses to noxious stimuli [12–15], yet their specificity for nociception over other sensory stimuli is not clear. Furthermore, it is currently unknown whether selective nociceptive coding in the ACC would be attained by single neuron representations or by coordinated dynamics of neuronal ensembles.

Patients suffering from chronic pain show elevated ACC activations associated with increased unpleasantness [1, 16–18]. In rodents, the transition to chronic pain is characterized by neuronal plasticity mechanisms in the ACC that contribute to nociceptive sensitization (Blom et al 2014, Santello & Nevian 2015; Bliss et al 2016). The overall nature of the reported changes point to increased excitability and stronger connectivity within the neuronal network. However, it is currently unknown how these plastic changes may affect network dynamics and coding capabilities of the ACC.

We chronically imaged ACC neuronal activity to understand how noxious sensory stimuli are encoded and distinguished from other salient or neutral stimuli, and what signatures of stimulus representation emerge in chronic pain. We took advantage of GRIN lenses to directly image from a population of ACC neurons with single-cell resolution, that otherwise were not in range for conventional cranial window imaging [19] (Figure 1A-B). Thus, this mesoscopic imaging technique allowed bridging the gap between global brain imaging studies using fMRI to single-cell electrophysiology. Animals under light isoflurane anaesthesia (0.5% - 0.9%) were presented with a battery of innocuous, aversive and noxious stimuli (Figure 1A-D). For each individual recording session, we also measured the spontaneous activity of neurons (Supplementary Figure 1A).

**Figure 1.**
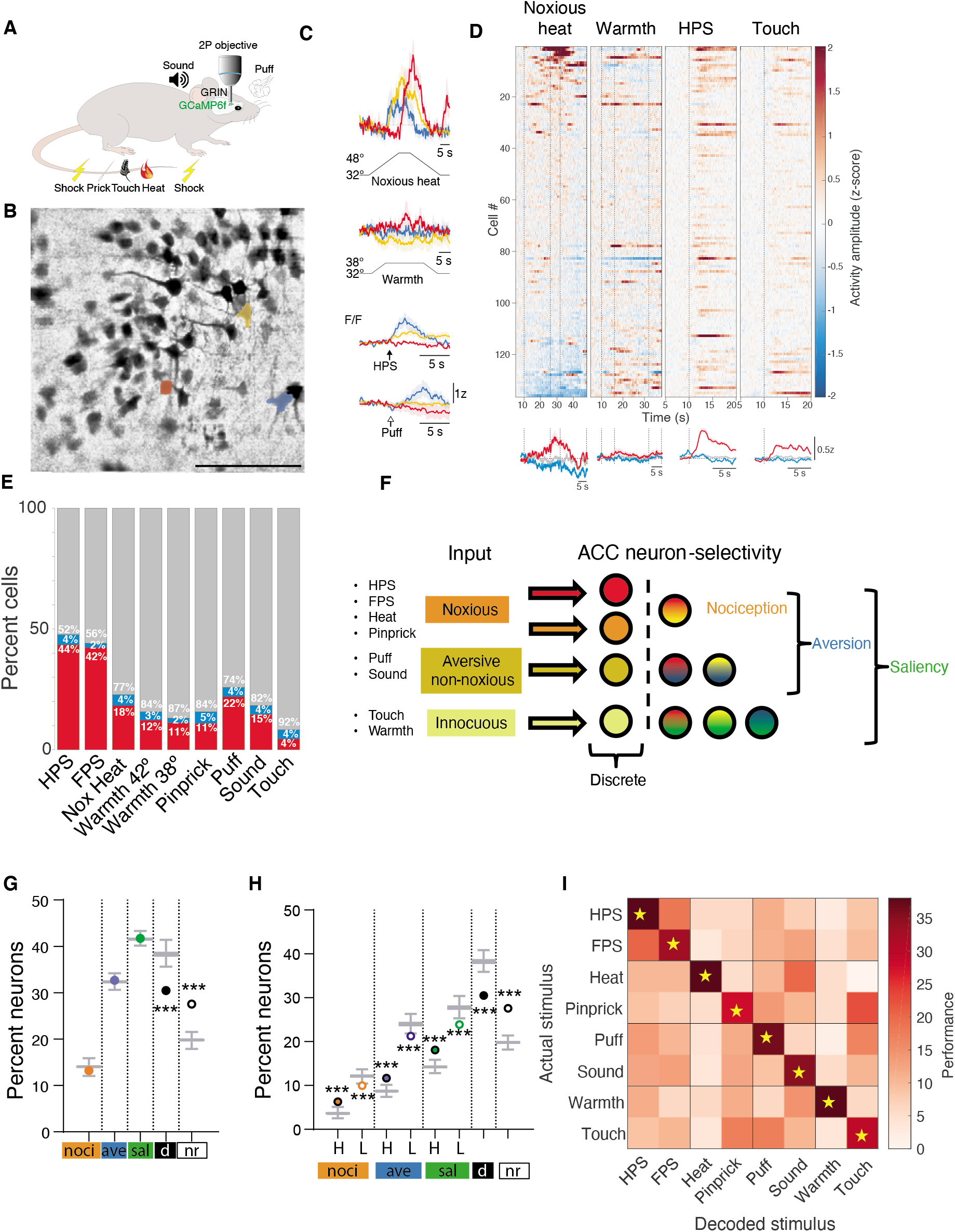
Representation of sensory concepts in the ACC. **(A)** ACC neuronal activity of GCAMP6f-expressing neurons was assessed through a GRIN lens under a two-photon microscope. Animals were kept in low isoflurane (0.5-0.9%) and different stimuli were delivered. Noxious heat, mild touch and pinprick were delivered exclusively to the contralateral hind-paw. Mild electric shock was presented contralaterally in both fore-paw and hind-paw. Air puff was delivered to the contralateral cheek-eye region. A loud tone was presented with equal volume to both ears. **(B)** Mean correlation image of a field of view (FOV) from a single session and animal denoting 3 individual neurons in L5 of the ACC. Scale bar: 100 μm. **(C)** Example of three neuronal activity responses (colour-coded from (B)) to heat, warmth, electric shock to the contralateral hind-paw and air puff. **(D)** Z-scored mean response activity across all trials imaged in a single session for the FOV shown in (A). Upper panel. neurons were aligned to the maximal response evoked by heat. Lower panel. Z-score activity response for either selective neurons (red, positively modulated; blue, negatively modulated) or non-responsive neurons to the corresponding stimulus. **(E)** Proportion of neurons responding to all stimuli delivered across 20 animals (1358 neurons). **(F)**. Schematic diagram of the strategy to categorize neurons according to its responsiveness to the stimulus set. This determines the concept to which a neuron is specific (i.e. nociception, aversion or saliency). **(G)** Percent of neurons that belong to each category described in (F). Box and whisker (mean and standard error) plots represent the chance distribution of 1000 simulated points. Circles are the actual observed values. *** p < 0.001 from chance based on Z scores. ‘noci’, ‘ave’ and ‘sal’ denote nociception, aversion and saliency-specific; ‘d’: discrete; ‘nr’: non-responsive. **(H)**. Same as (G) with cells further subdivided in high (H) and low (L) concept responders, based on their number of specific stimuli. *** p < 0.001. **(I)** Performance of a logistic regression decoder that classified neuronal activities for all stimuli presented. The performance of on-diagonal (actual stimulus) values was compared to all off-diagonals (incorrect stimuli) and normalized to 100%. Yellow stars indicate the correct classification (p < 0.05) of the actual stimulus over off-diagonal stimuli within the same row (Wilcoxon sign-rank).

We first investigated whether the ACC possessed individual dedicated nociceptive neurons. We developed machine learning algorithms to detect stimulus-evoked neuronal responses. We classified then neurons being responsive, non-responsive or inhibited based on the activity pattern after stimulus onset (Supplementary Figure 2). Overall, across sessions, sensory stimuli elicited significant Ca^2+^ responses in 72.42% of imaged ACC neurons (1358 neurons; 67.9 ± 4.4 neurons per session per animal). We saw that mild electric shock applied to the hind-paw (HPS) recruited the highest number of neurons, whereas innocuous mild touch activated the smallest subset of neurons (Figure 1E). A large portion of the activated neurons responded to one stimulus (~42% of total neurons), whereas ~58% were multi-modal and responded to two or more stimuli (Supplementary Figure 1B). For all stimuli tested, individual neuronal response reliability was typically low (18.87 % ± 0.16; Supplementary Figure 1C).

To identify nociception-specific neurons under our experimental conditions, we classified all cells in five groups as follows: i) non responsive; ii) discrete: responsive to only one out of all stimuli (individual discrete neurons shown in Supplementary Figure 1D); iii) nociception-specific: activated exclusively by two or more noxious stimuli; iv) aversion-specific: responding exclusively to two or more aversive (noxious or not) stimuli; v) saliency-specific: activated by two or more stimuli (Figure 1F). The logic of this classification lies in identifying the common concept to all stimuli to which a given neuron responds. This way, the nociceptive ensemble will be composed of cells only activated by stimuli that share the property of engaging the nociceptive system, and not to other aversive or salient stimuli. Accordingly, discrete neurons responding to a single noxious stimulus are excluded from the ensemble because it is not clear what quality of the event is encoded. After sorting all neurons according to these criteria, we compared the proportions observed for each category to a chance distribution obtained by bootstrapping 1000 times the identity of the activated cells. We observed that the number of neurons encoding for nociception, aversion or saliency was not different from what was expected by chance (Figure 1G). However, we found less discrete and more non-responsive cells than expected by chance, suggesting that the ACC might encode generalized stimulus qualities (i.e. concepts) rather than specific representations. To further characterize this hypothesis, we divided nociception, aversion and saliency-specific cells into low (L) and high (H) concept responders, according to the number of stimuli by which they were activated (L: only 2; H: 3 or more). We found that the number of H neurons from all three classes was greater than chance (Figure 1H), whereas the amount of L cells was smaller than expected (Figure 1H, Supplementary Figure 1E). Altogether, this data indicate that ACC neuronal networks represent noxious events by means of a distributed neuronal network that includes highly nociception-, aversion- and saliency-specific concept responding neurons (~17% of recorded cells), which combined comprise 64% of all responses to nociceptive stimuli.

The combination of neurons representing individual stimuli with concept-selective cells may be suitable to discriminate nociceptive and non-noxious sensory events with different valences, intensities, and modalities. To assess this, we performed machine learning classification methods in order to evaluate how the entire neuronal population can distinguish a given stimulus. We found that the ACC can effectively discriminate between stimuli (Figure 1I).

Overall, these results provide evidence for a robust representation of the sensory space in ACC neuronal ensembles described by its characteristic pattern of activity. Moreover, our data suggests that nociception-specificity could be attained by of a core of ACC neurons that encode the different concepts underlying the noxious experience, like nociception, aversion and saliency.

The encoding of stimulus features in individual neurons is generally stable over time in primary sensory cortices [20–25]. However, the stimulus representation on the cellular level tends to continuously reconfigure in associative areas despite stable behavioral performance [25, 26]. It is currently unknown how the temporal dynamics of nociceptive representation in higher cortical association areas evolve over time on the cellular level. Resolving this question will help to understand if nociception-specificity is achieved through hard-wired, stable and dedicated neurons or by flexible and dynamic neuronal ensembles. We therefore evaluated the neuronal representational changes over time in up to three consecutive sessions separated by a week in naive animals (Figure 2A, Supplementary Figure 3A). The spontaneous activity of individual neurons correlated across sessions (Figure Supplementary 3B). However, their responses to the different stimuli were highly variable from one session to another, showing considerable reconfiguration of single cell tuning (Figure 2A, Supplementary Figure 3C). To assess how these dynamics impacts on the population code, we focused on a selection of stimuli that showed the highest proportion of activated neurons, i.e. heat, HPS and puff. We identified neurons that gained responsiveness, neurons that lost responsiveness, neurons that remained stable and neurons that changed the valence of responsiveness (swap neurons) (Figure 2B). We observed that a large portion of neurons changed their identity to a given stimulus and that only a smaller fraction was stable. Nevertheless, the total number of neurons in a stimulus ensemble remained constant as there was an apparent homeostatic balance of equal amounts of neurons gaining and losing responsiveness (Figure 2B, Supplementary Figure 3D, net variability = −0.77 ± 5.8). Next we assessed the stability of response selectivity and reliability, two attributes which carry information about the quality of the stimulus representation. We compared the magnitude of gain versus lost response selectivity and reliability across sessions. For all the evaluated stimuli, neither of the two parameters changed (Figure 2C, Supplementary Figure 3E). Thus, despite the identity of most neurons representing individual stimuli changes over time, sensory stimulus representation at the population level remains stable in quantity (proportion) and quality (selectivity and reliability). In this way, the brain can achieve stability with a high degree of flexibility [26]. If this is true, we expected that information contained in stimulus-evoked neuronal activity might allow for generalizability of the neuronal code across sessions. To test this hypothesis, we trained a decoder to classify stimulus-evoked patterns in a given session, and tested on the activity patterns of the same neurons recorded in the subsequent session. We found that accuracy dropped over sessions (Figure 2D), because of the changes in single-cell identity and response activity between days. Nevertheless, performance was still considerable (0.70 ± 0.018), demonstrating a partially stable representation of the sensory code in ACC neuronal ensembles. We next examined the temporal stability of nociception, aversion and saliency-specific H neurons (i.e. highly concept-responding neurons) between two consecutive sessions. First, we observed that despite subtle changes in the proportion of neurons that belong to each category, significant representation above chance levels remained stable over time (Figure 2E Figure Supplementary 3F). Then, we analyzed whether the identity of concept-specific neurons changed from one session to the next. We found a significant proportion of H neurons that kept their specificity for nociception, aversion and saliency (9.75%, 27% and 46%, respectively). For all cases, the number of cells was higher than expected by chance and together with stable non-responding cells add up to ~20% of all recorded ACC neurons (Figure 2F, Supplementary Figure 3G). Our data identified for the first time a core of ACC neurons showing perdurable specificity for abstract concepts such as nociception, aversion and saliency. This, together with stable population encoding parameters, may provide the ACC with the ability to keep sensory representation constant in the presence of continuous change [26, 27].

**Figure 2.**
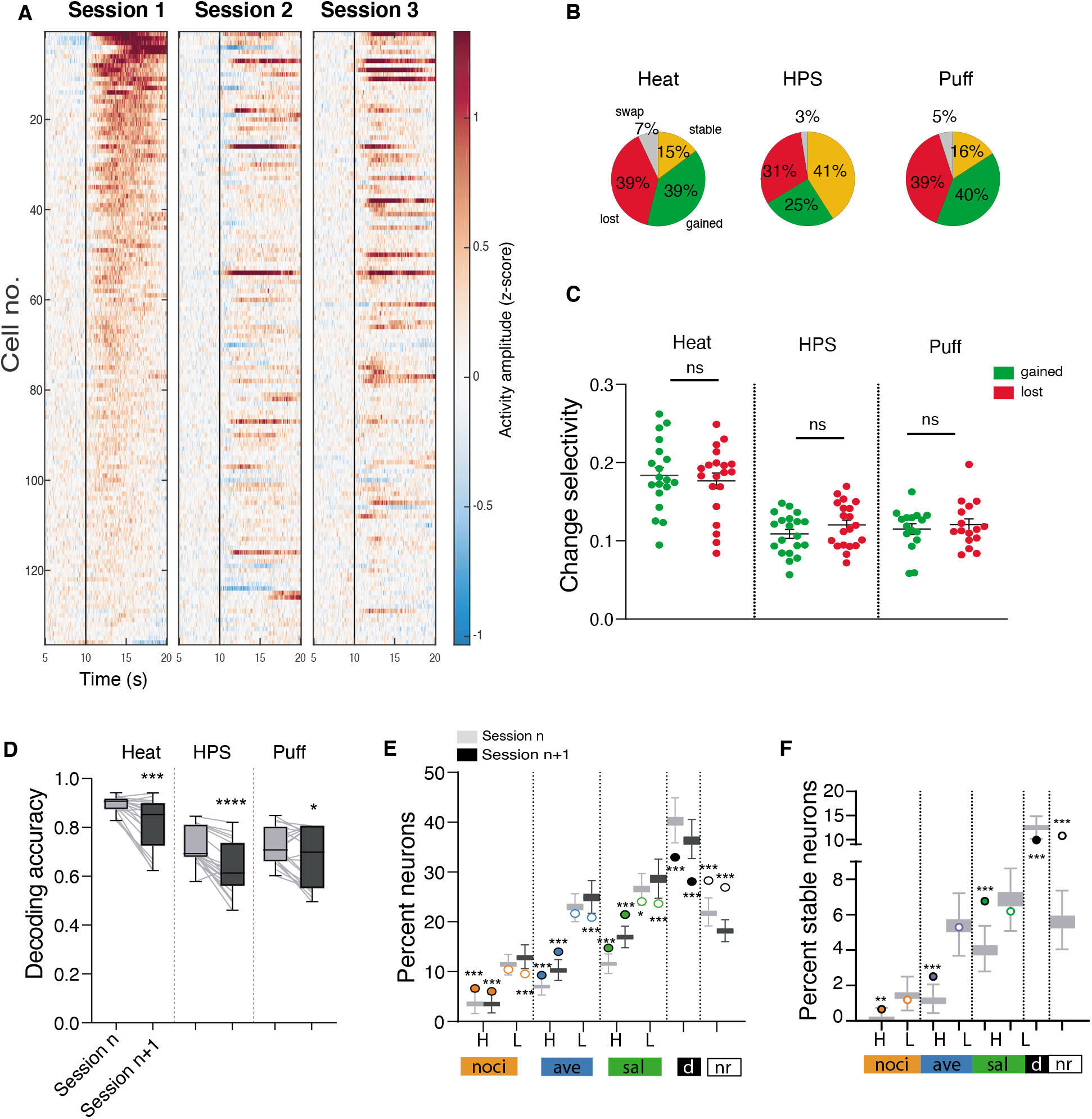
Temporal stability of stimulus representation in the ACC. **(A)** Time-lapse stimulus response activity (z-score) for HPS across three consecutive sessions, sorted to the neuron with the highest stimulus-evoked activity in session one. **(B)** Pie charts depicting the proportion of neurons that either gained, lost, changed modulation (from inhibited to activated, or vice versa - swap), or remained stable across sessions for heat, HPS and puff. **(C)** Net change in selectivity over time per animal calculated as the difference of gained and lost neurons, divided by the number of selective neurons in two consecutive sessions. ns: p > 0.05, unpaired t-test, n = 19 - 20 animals. **(D)** Generalizability, measured as accuracy difference between two consecutive sessions for heat, HPS and puff. * p < 0.05, *** p <0.01, **** p < 0.001, paired t-test, n = 20 animals. **(E)** Percent of neurons that belong to each category in two successive sessions. Box and whiskers represent the chance distribution and circles are the actual observed values. * p <0.05, *** p < 0.001 from chance based on z-scores. H and L refer to high and low concept responsive cells. ‘noci’, ‘ave’ and ‘sal’ denote nociception, aversion and saliency-specific; ‘d’: discrete; ‘nr’: non-responsive. **(F)**. Percent of neurons that conserve category-specificity in two successive sessions. Box and whiskers represent the chance distribution and circles are the actual observed values. ** p <0.01, *** p < 0.001 from based on Z scores. Labels are the same as (E).

ACC neuronal circuits undergo substantial synaptic and intrinsic modifications during the development of chronic pain [28–30], which are believed to play a central role in its symptomatology [29, 30]. Therefore, we aimed to study how stimulus representation and neuronal dynamics were affected in neuropathic pain. We used the chronic constriction injury (CCI) of the sciatic nerve as a model for chronic pain (Figure 3A). One group of animals underwent CCI, whereas the other group underwent sham surgery. We performed calcium imaging of the same field of view, tracking the same set of cells longitudinally before and after surgery (Figure 3B). From a total of 20 animals (8 sham and 12 neuropathic) we were able to follow 1358 neurons. Throughout the development of chronic neuropathic pain, the reconfiguration of individual sensory stimulus representations remained stable, measured by the rate of gained/lost cells and stimulus selectivity (Figure Supplementary 4 A-B). Moreover, nerve injury did not significantly alter spontaneous activity of ACC neurons (Figure Supplementary 4C-D) nor the response magnitude of neurons activated by nociceptive, aversive or innocuous stimuli (Figure 3C-D). However, while neuropathic pain did not alter the proportion of neurons engaged by nociceptive and aversive stimuli, two weeks after sciatic nerve surgery, light touch recruited a higher number of neurons in CCI mice (Figure 3E-F). Recently, neuropathic pain-related allodynia was associated with an expansion of innocuous stimulus representation toward nociception-responsive neurons in the basolateral amygdala [31]. To test if a similar phenomenon occurred in the ACC, we measured the overlap between touch and nociceptive responsive neurons. We found that compared to naïve conditions, there was a significant increase in the proportion of cells activated simultaneously by touch and noxious heat, and by touch and shock in neuropathic animals but not in sham operated animals (Figure 3E-G). Then, we evaluated how neuropathic pain impacted the structure of nociception, aversion and saliency-specific ensembles. Contrary to sham animals, where the representation of H selective cells from all categories remained above chance levels after surgery, nociception and aversion-selective H-concept neurons were impaired in neuropathic animals (Figure 3H, Supplementary Figure 4E). Conversely, the number of saliency-selective H cells remained higher than expected by chance in neuropathic mice, likely due to the expansion of touch-responsive cells to the nociceptive ensemble. In agreement with this, the contribution of innocuous stimuli-responsive cells to the H saliency-specific ensemble was larger after sciatic nerve injury (Figure 3I).

**Figure 3.**
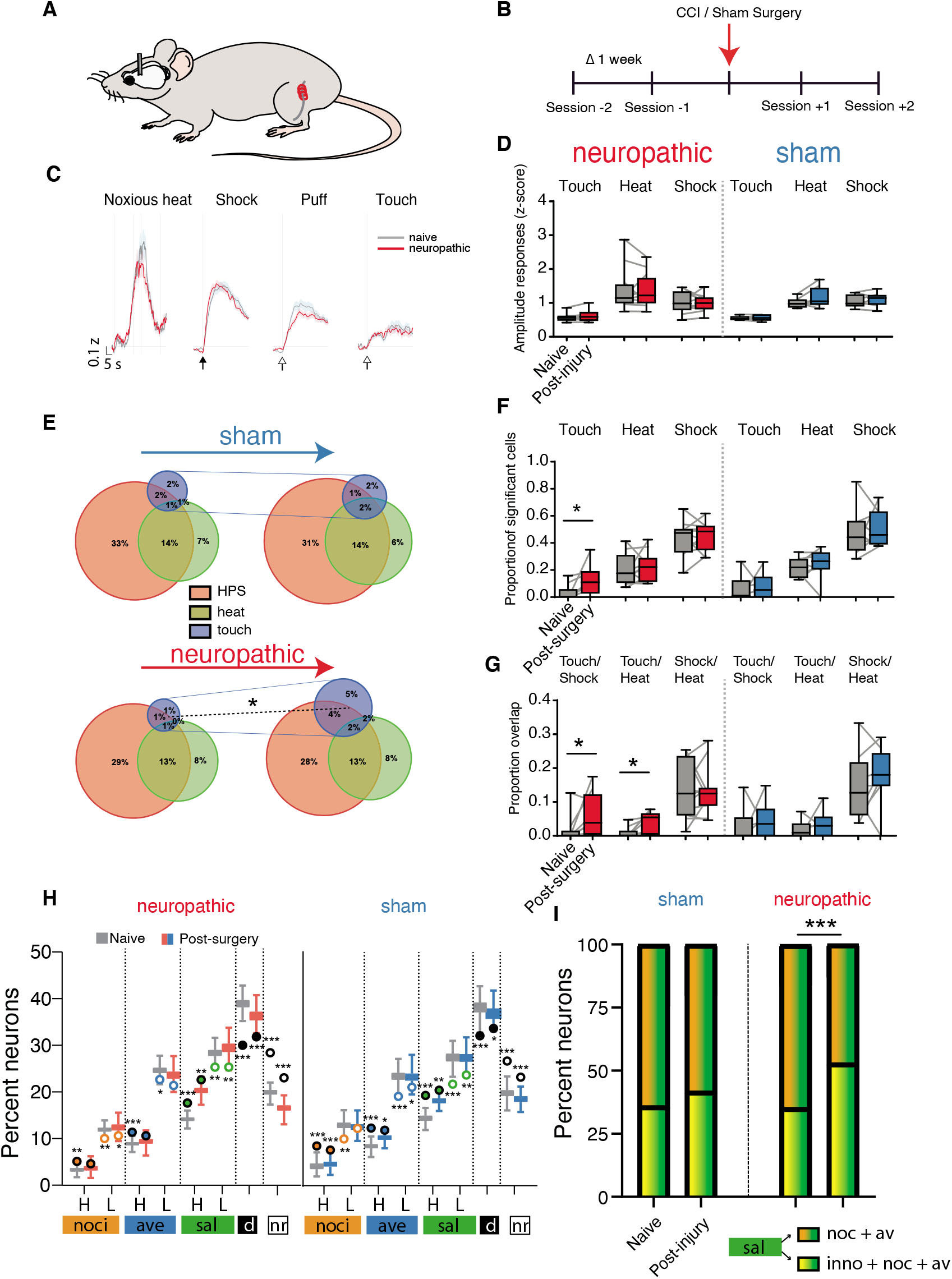
Temporal dynamics of sensory representation in neuropathic pain. **(A)** Schematics of the chronic constriction injury model of neuropathic pain. The sciatic nerve of the left leg was ligated. Three ligatures were loosely tied around the nerve. **(B)** Time schematics of the recording sessions. Imaging was performed at a one week interval. **(C)** Averaged z-score traces of stimulus response activity to heat, shock, puff and touch before (naive, grey) and after CCI (neuropathic, red). Thick line: mean, shadow: SEM. **(D)** Amplitude response per animal pre- and post-surgery. p > 0.05, Wilcoxon rank-sum. Naive-neuropathic, n = 8-12. Naive - sham, n = 6-8. **(E)** Venn diagrams depicting the change of the proportion of responding cells in naive and post-surgery (sham and neuropathic) conditions. The number of responding neurons is significantly larger after surgery only for the neuropathic group * p < 0.05. Wilcoxon rank-sum. **(F)** Proportion of significant neurons modulated by touch, heat and shock before and after nerve injury. * p < 0.05. Wilcoxon rank-sum. Naive-neuropathic, n = 8-12. Naive - sham, n = 6-8. **(G)** Proportion of neurons with overlap in selectivity. * p < 0.05. Wilcoxon rank-sum, Naive-neuropathic, n = 8-12. Naive - sham, n = 6-8. **(H)** Percent of neurons that belong to each category in naïve conditions and after sciatic nerve surgery. Box and whiskers represent the chance distribution and circles are the actual observed values. * p <0.05, *** p < 0.001 from chance based on z-scores. H and L refer to high and low concept responsive cells. ‘noci’, ‘ave’ and ‘sal’ denote nociception, aversion and saliency-specific; ‘d’: discrete; ‘nr’: non-responsive. **(I)** Relative contribution of aversion-specific and innocuous stimuli responsive cells to the H saliency-specific ensemble. *** p < 0.001. Fischer test for proportions.

Taken together, these data demonstrate that neuropathic pain affects sensory stimulus representation in two ways. First, innocuous stimuli engage a larger fraction of nociception-responsive cells. Second, nociception and aversion encoding by dedicated neurons is disrupted, probably reflecting changes in connectivity patterns of ACC microcircuits [28]. Based on this, we hypothesized that the emergence of the allodynic neuronal phenotype might not only depend on the expansion of the innocuous ensemble, but also on the inability to differentiate innocuous from nociceptive stimuli.

To gain insight into this, we aimed to characterize the relationship between neuronal selectivity for nociceptive and innocuous stimuli. We first clustered neurons based on selectivity between pairs of stimuli. We noticed that the distance in the selectivity space between nociceptive and innocuous stimuli is largely reduced in neurons from neuropathic animals (Figure 4A, Figure supplementary 5A). This suggests that the selectivity tuning for innocuous stimuli resembles those of noxious stimuli only in neuropathic animals. This may therefore lead to a less distinguishable stimulus representation. To corroborate this hypothesis, we performed k-means clustering analysis of the neuronal selectivity for categories of stimuli. We observed that the distance in the multidimensional space between clusters of neuronal selectivities for noxious and innocuous stimuli lay significantly closer in neuropathic animals compared to sham animals (Figure 4B). However, the distance between aversive and innocuous as well as noxious to aversive was not changed (Figure supplementary 5B). This further corroborates the idea that the allodynic phenotype -a hallmark of chronic pain - may not only depend on the expansion of neuronal ensembles, but also on the increased similarity of intrinsic stimulus responses between noxious and innocuous stimuli. Based on these data, it is therefore tempting to hypothesize that neuropathic pain may lead to a misclassification of innocuous stimuli as noxious. To test this idea, we trained a classifier to assess the correct decoding of individual stimuli in neuropathic animals as well as in sham animals. First, whereas all stimuli were properly discriminated in sham animals, the discrimination of two nociceptive stimuli, noxious heat and pinprick, was impaired in CCI mice (Figure 4C). Interestingly, we noticed an increase of false classification of innocuous stimuli as noxious stimuli (Figure 4C, Fig Supplementary 4C) in neuropathic pain animals, compared to sham animals. This suggests that the underlying quality of the neuronal representation of innocuous stimulus resembles that of noxious-evoked neuronal activity.

**Figure 4.**
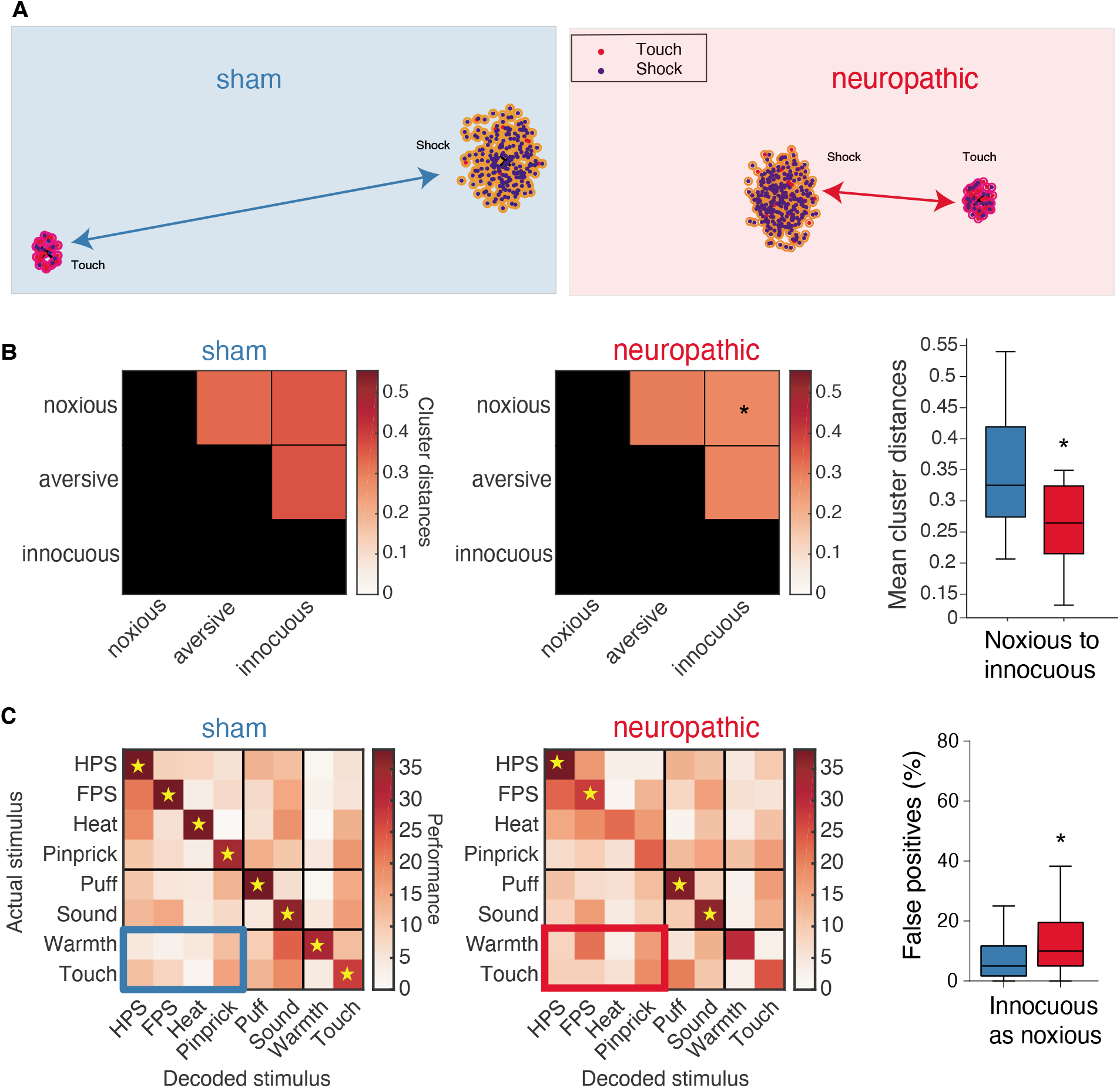
Neuropathic pain alters cortical classification of stimuli. **(A)** Clustering of neurons in the selectivity map using the UMAP algorithm. Colors surrounding each dot correspond to the classification by k-means. Colors inside each circle corresponds to the favorable selectivity of each neuron. **(B)** k-means cluster analysis for categories of stimuli, noxious, aversive or innocuous. * p < 0.05, neuropathic n = 31 clusters, sham n = 16 clusters. **(C)** Percent of false positives of the logistic regression classification algorithm for each individual stimulus for sham and neuropathic animals. Yellow stars indicate the correct classification (p < 0.05) of the actual stimulus over off-diagonal stimuli within the same raw (Wilcoxon sign-rank). Blue and red box indicates the falsely classified innocuous stimuli (warmth and touch) as noxious stimuli (HPS, FPS, heat and pinprick). Neuropathic, n = 55. Sham, n = 44 combinations. * p < 0.05. Wilcoxon rank-sum.

Thus, our data demonstrates changes in the ACC population coding in neuropathic pain due to a defective representation of nociceptive and aversive attributes by dedicated neurons, to an expansion of the neuronal ensemble recruited by innocuous stimuli, and to alterations in the intrinsic features of neuronal activity at a network level. Together, this accumulates to an aberrant encoding of the nociceptive sensory space in the ACC during chronic pain.

## Acknowledgements

We thank Natalie Nevian for excellent technical assistance. This work was supported by the Swiss National Science Foundation (T.N., grant 159872; T.N. and F.K. 173486) and the European Research Council (T.N., grant 682905).

## Author contributions

M.A.A., F.K., T.N. designed the experiments and wrote the manuscript. M.A.A. and F.K. performed the experiments. M.A.A., F.K., P.D.L., analyzed data. All authors contributed to the manuscript.

## Supplementary figure legends

**Supplementary Figure 1.**
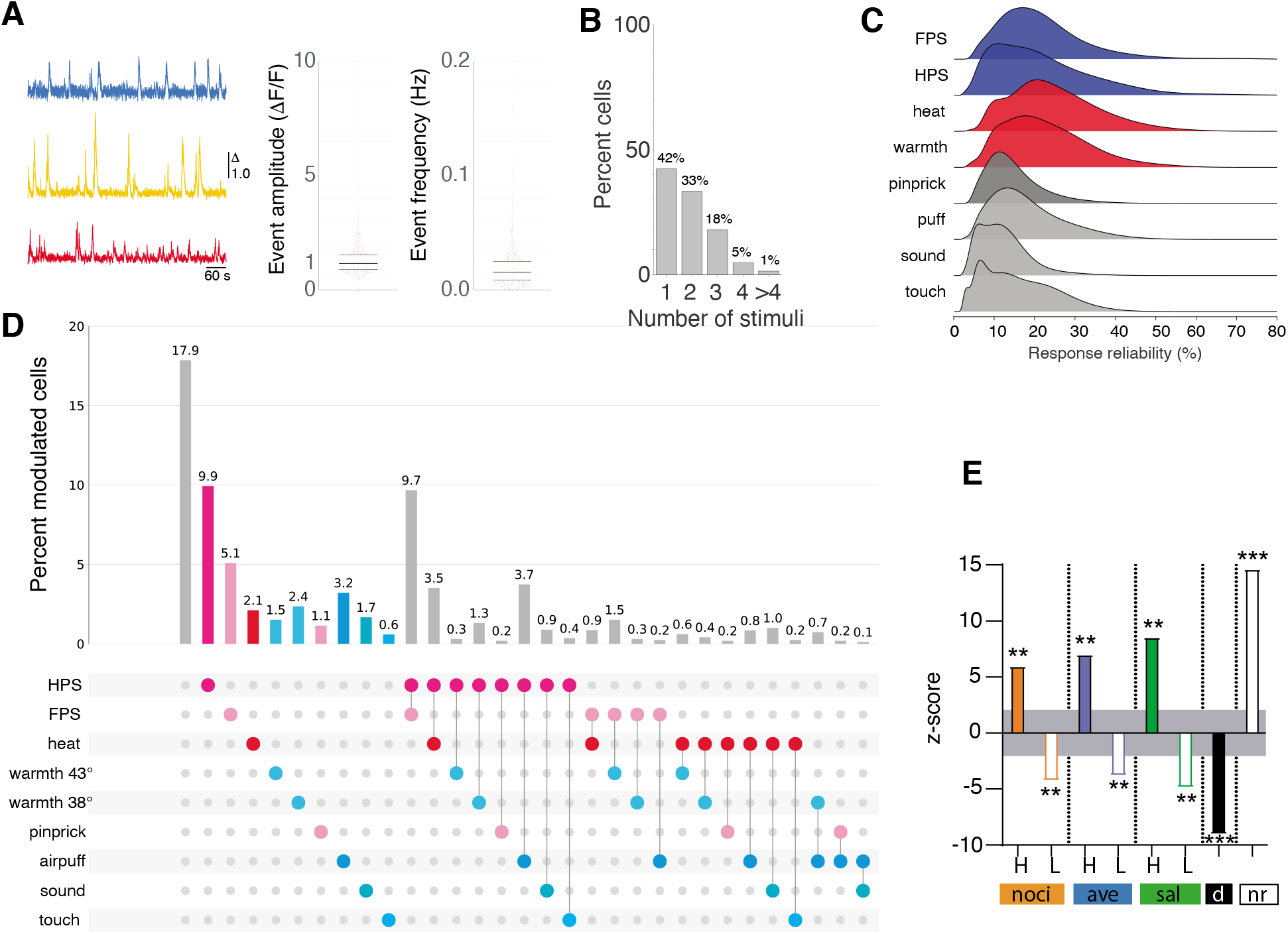
Associated data to Figure 1. **(A)** Left, traces of spontaneous activity of individual neurons from Figure 1B-C. Right, scatter plot for all neurons analyzed. n = 1358 neurons. **(B)** Proportion of unimodal (neurons responding to only one stimulus) and multimodal (neurons responding to more than two stimuli). **(C)** Density plot depicting the response reliability of neurons to the different stimuli. **(D)** Combinatorial of stimuli and the respective proportion of neurons specific to a combination (up to two stimuli). **(E)** Z-score calculated for real observed percent of neurons relative to the chance distribution parameters belonging to each category described in Figure 1F. Grey area indicates the significance threshold of ± 2 standard deviations. ** p <0.01, *** p < 0.001. H and L refer to high and low category responsive cells. ‘noci’, ‘ave’ and ‘sal’ denote nociception, aversion and saliency-specific; ‘d’: discrete; ‘nr’: non-responsive.

**Supplementary Figure 2.**
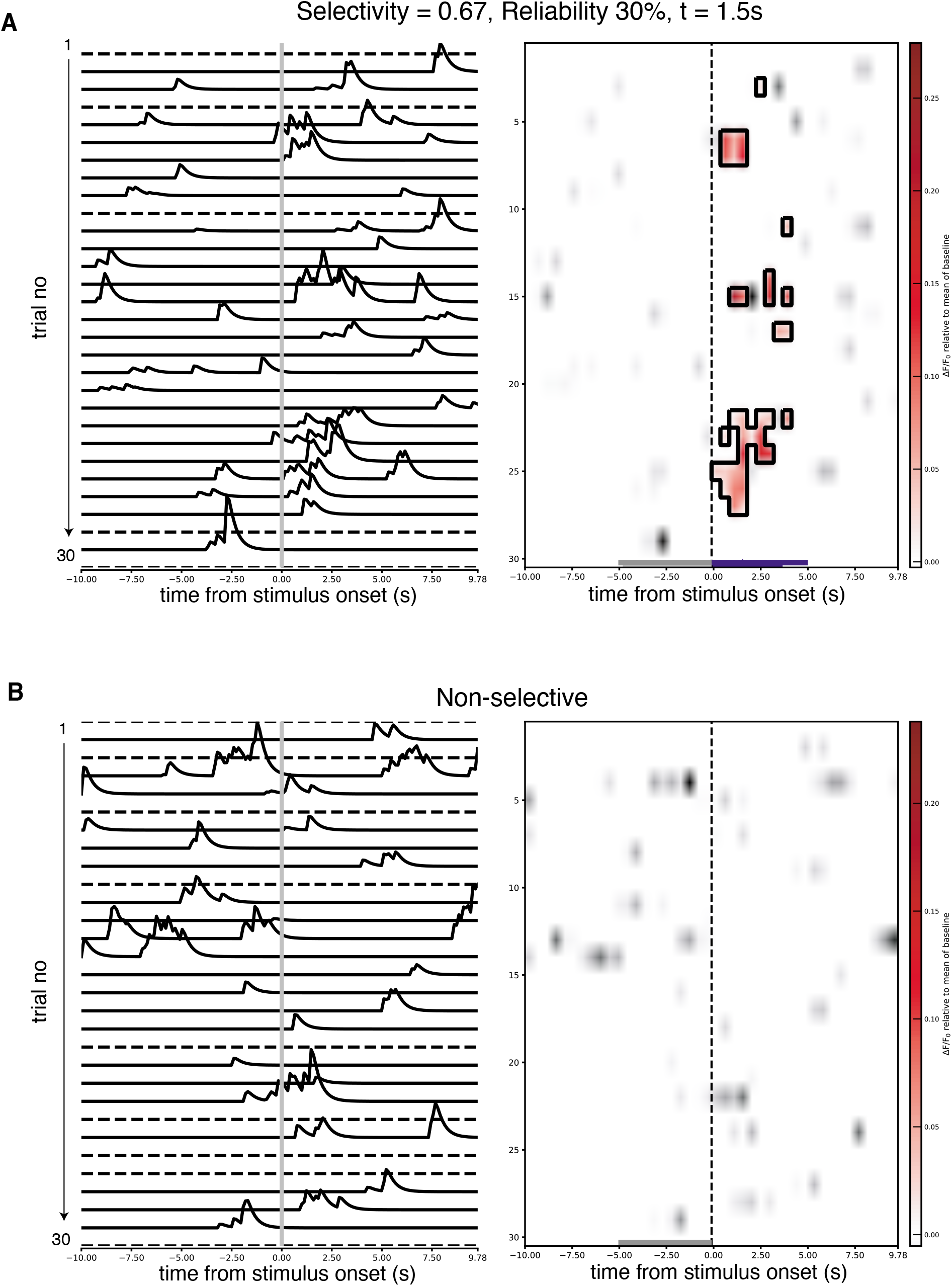
Classification of selective neurons. **(A)** Example of a selective neuron to HPS. Our classification method detects single trial activations, respective to the baseline activity before stimulus onset. Left, denoised traces. Right, activation detection. Black squares, significant activation periods within the 5 second-window (purple line), compared to baseline (grey line). **(B)** Same as in (A), but for a non-selective neuron. Even though the neuron shows activity after the stimulus onset, this does not pass the statistical significance to classify it as responsive given the prior activity.

**Supplementary Figure 3.**
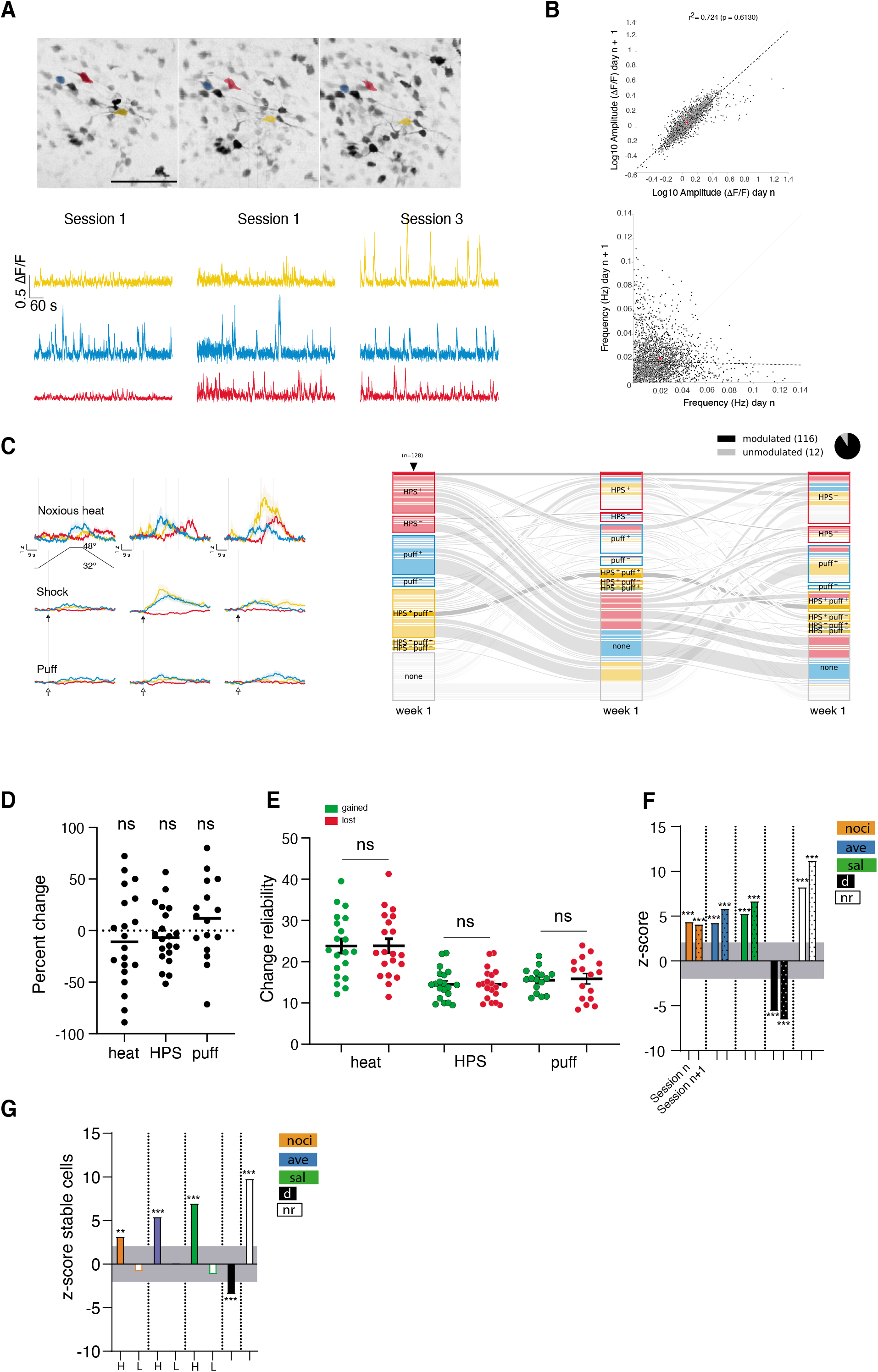
Associated data to Figure 2. **(A)** Top panels. Single field of view (FOV) tracked over sessions. Scale bar: 100 μm. Lower panel. Spontaneous activity of colored neurons in top panels. **(B)** Spontaneous activity correlation across sessions. One session (n) is compared to a subsequent session (n + 1). Note the high correlation of calcium event amplitudes across sessions. **(C)** Z-scored mean response activity across sessions of the neurons shown in (A), denoting the change in stimulus-evoked activity. Line, average; shadow, SEM. Right panel. Alluvial plot depicting the change in neuronal selectivity over time for HPS and puff. ‘None’ indicates a population of neurons not responding to either stimulus. Each individual line represents a single neuron tracked over time. (+) indicates positively modulated neurons and (-) denotes negative modulated neurons to their respective stimulus. Arrowhead indicates the initial tracking session where colors relate to a given selectivity. Colors in other columns represent the respective initial selectivity (at day 1). Pie chart corresponds to the total number of cells modulated within these three sessions. **(D)** Net change in number of neurons that gained or lost selectivity across sessions per animal relative to the number of selective neurons in both sessions for heat, HPS and puff. ns: p > 0.05, n = 16-20, one-sample t-test against zero. **(E)** Reliability change, measured as the response probability, in neurons that gained or lost selectivity. ns: p > 0.05, n = 16-20, unpaired t-test. **(F)**. Z-score calculated for real observed percent of neurons relative to the chance distribution parameters that belong to each category described in Figure 2H. Grey area indicates the significance threshold of ± 2 standard deviations. *** p < 0.001. ‘noci’, ‘ave’ and ‘sal’ denote nociception, aversion and saliency-specific; ‘d’: discrete; ‘nr’: non-responsive. **(G)** Same as in (F) but z-score for stable neurons specific for each category. ** p <0.01, *** p < 0.001.

**Supplementary Figure 4.**
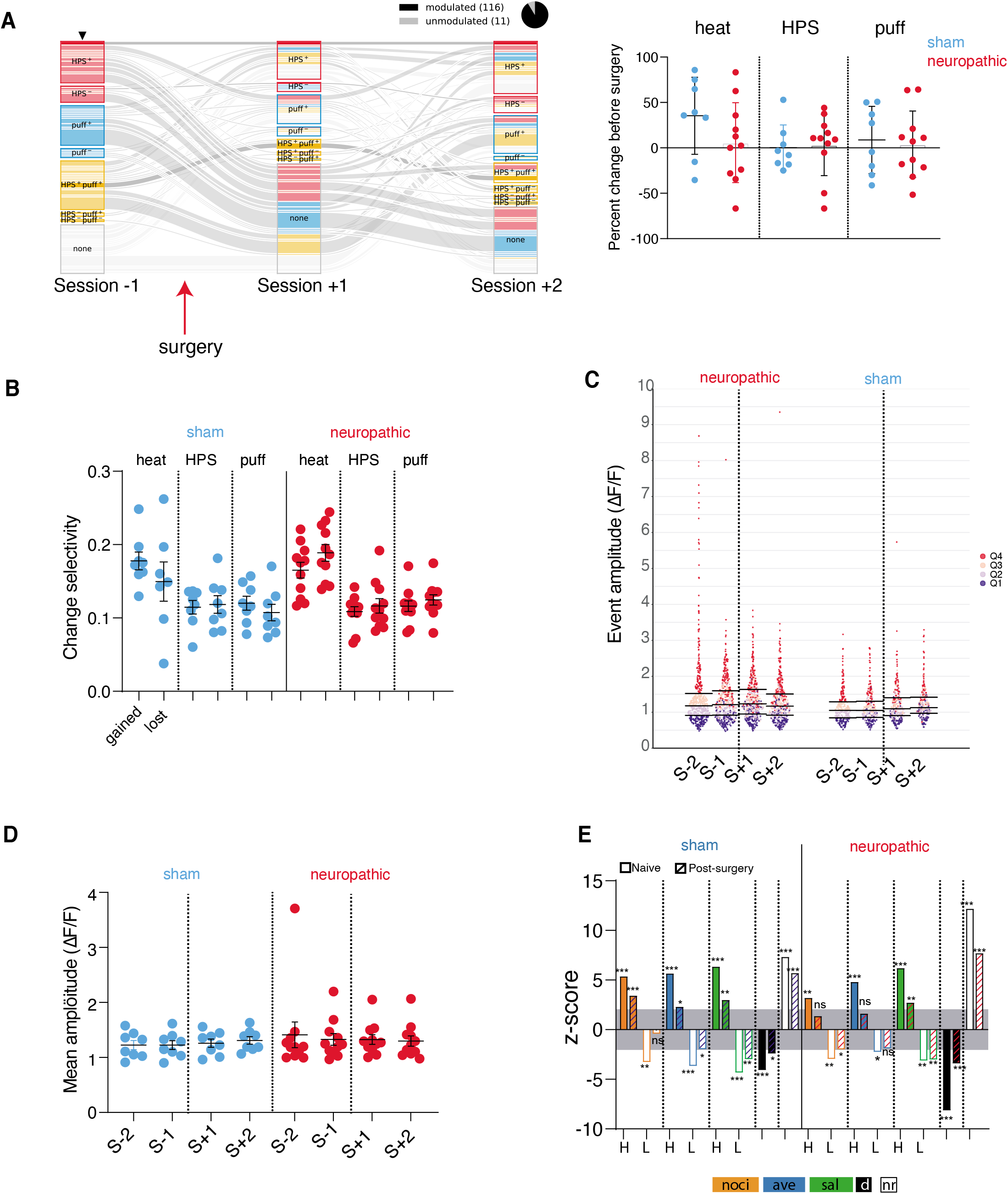
Associated data to Figure 3. **(A)** Left panel. Alluvial plot depicting the change in neuronal selectivity over time for HPS and puff. ‘None’ indicates a population of neurons not responding to either stimuli. Each individual line represents a single neuron tracked over time. (+) indicates positively modulated neurons and (-) denotes negative modulated neurons to their respective stimulus. Arrowhead indicates the initial tracking session where colors relate to a given selectivity. Colors in other columns represent the respective initial selectivity (in Session −1). Pie chart corresponds to the total number of cells modulated within these three sessions. Right panel. Scatter plot depicting the net percent change in number of gained and lost neurons over the total number of selective neurons for heat, HPS and puff, at Session +2, relative to Session −1 (before). p > 0.05 for all data sets, one-sample t-test against zero, sham n = 8, neuropathic n = 11-12 animals. **(B)** Selectivity change across sessions after surgery compared with before surgery for sham animals and neuropathic animals. p > 0.05 for all data sets, paired t-test per stimulus, sham n = 8, neuropathic n = 11-12 animals. **(C)** Scatter plot of all neurons recorded across four sessions (two sessions before and two sessions after surgery) from sham and neuropathic animals. Data sets are divided into quartiles, sorted at Session-2 (S-2). Note the stability of neurons present in the same quartile across sessions. Neuropathic n =779, sham n = 473 identified neurons. **(D)** Mean amplitude per animals across sessions. Event amplitudes remain stable across time. Sham = p > 0.05, F (2.147, 15.03) = 3.338, One-Way Repeated Measures ANOVA, n = 8. Neuropathic = p > 0.05, F (1.053, 10.53) = 0.3895, One-Way Repeated Measures ANOVA, n = 11. **(E)** Z-score calculated for real observed percent of neurons relative to the chance distribution parameters that belong to each category described in Figure 3H. Grey area indicates the significance threshold of ± 2 standard deviations.* p <0.05, ** p <0.01, *** p < 0.001. H and L refer to high and low concept responsive cells. ‘noci’, ‘ave’ and ‘sal’ denote nociception, aversion and saliency-specific; ‘d’: discrete; ‘nr’: non-responsive.

**Supplementary Figure 5.**
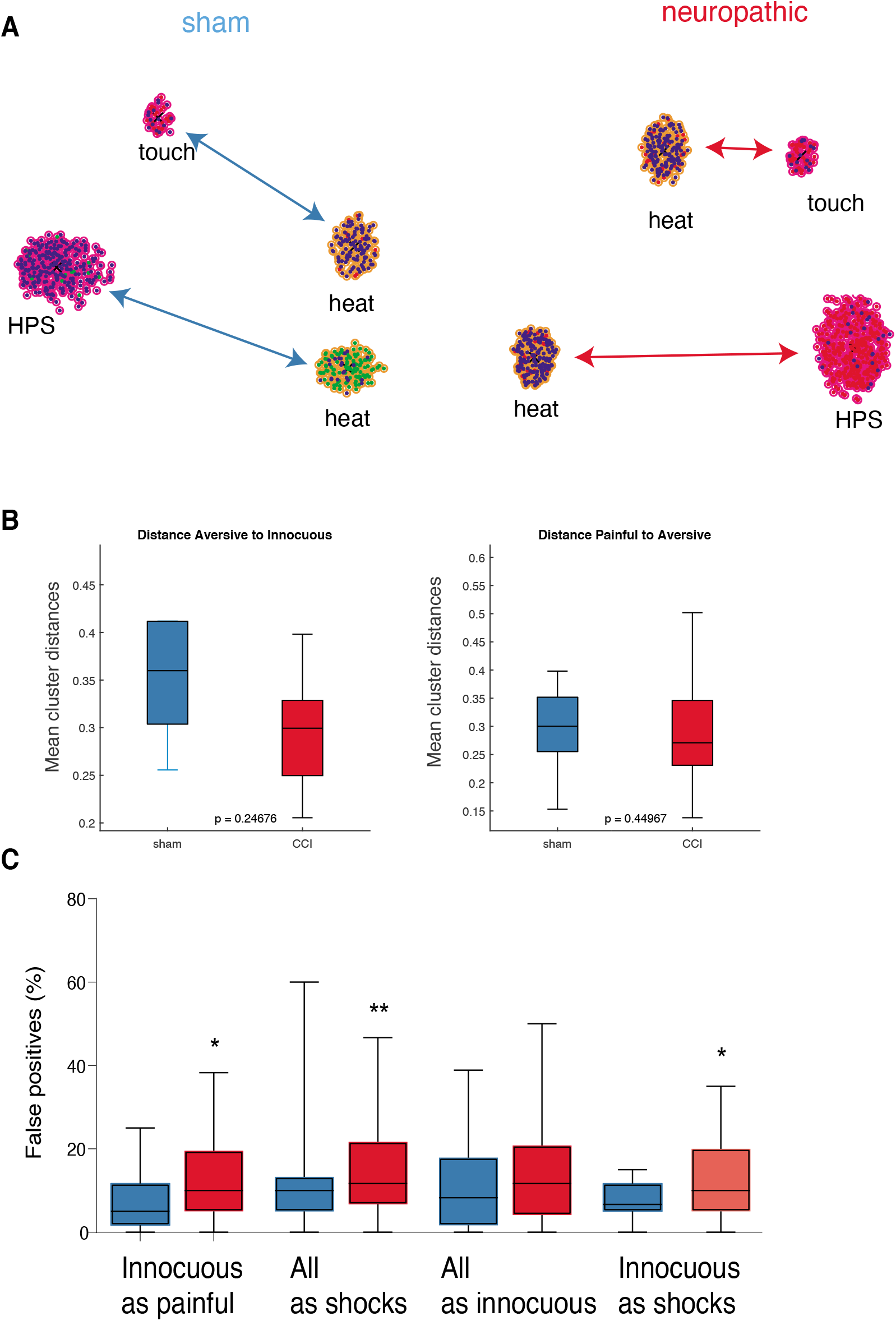
Associated data to Figure 4. **(A)** Clustering of neurons in the selectivity map using the UMAP algorithm. Colors surrounding each dot correspond to the classification by k-means. Colors inside each circle corresponds to the favorable selectivity of each neuron. Note the distance is noticeable shorter between innocuous (touch) and noxious (heat) only in neurons from neuropathic animals, whereas the distance between two noxious stimuli (HPS-heat) remains constant. **(B)** K-means cluster analysis for categories of stimuli. Left panel. Distances between aversive and innocuous stimuli are not significantly different. p > 0.05, Wilcoxon sum-rank. Neuropathic n = 12 clusters, sham n = 5 clusters. Distances between painful and aversive stimuli are not significantly different. p > 0.05, Wilcoxon sum-rank, neuropathic n = 43 clusters, sham n = 20 clusters. **(C)** Percent falsepositive rate of logistic regression classification of stimuli in categories. * p < 0.05, ** p < 0.01. Innocuous as painful, Neuropathic, n = 55. Sham, n = 44 combinations. * p < 0.05. Wilcoxon rank-sum. All as shocks, Neuropathic, n = 98. Sham, n = 89 combinations. ** p < 0.01. Wilcoxon rank-sum. All as innocuous, Neuropathic, n = 56. Sham, n = 45 combinations. p > 0.05. Wilcoxon rank-sum. Innocuous as shocks, Neuropathic, n = 28. Sham, n = 23 combinations. * p < 0.05. Wilcoxon sum-rank.

## Methods & statistical analysis

### 1. Animals

All experiments were conducted in accordance to the rules of the veterinary office of the canton of Bern, Switzerland. Male C57/BL6 mice (Central Animal Facility of the University of Bern or Janvier Labs, France) were housed in groups of 4-5 per cage and maintained on a 12-hour light/dark cycle in a temperature-controlled environment with ad libitum access to food and water.

Surgeries

#### 1.1. Virus injections

Viral vectors used to express the genetically-encoded calcium indicator GCaMP6f (AAV1.Syn.GCaMP6f.WPRE.SV40; AAV1.CAMKII.GCaMP6f.WPRE.SV40) were obtained from Penn Vector Core, Gene Therapy Program, Perelman School of Medicine, University of Pennsylvania, US. Viral vectors were aliquoted and stored at −80°C until the day of surgery, when were diluted 1:5 in PBS before use.

All surgeries were conducted under aseptic conditions with glass bead sterilized surgical tools (Fine Science Tools). Adult mice (~6 weeks old) were anaesthetized with isoflurane (3–4% induction, 1–2% maintenance) vaporized with 100% oxygen and placed in a stereotaxic frame (Stoelting, USA). Eye ointment (Pharma Medica AG, Rogwil, Switzerland) was applied and body temperature was monitored and maintained at 35-37°C using a heating pad (brand). After cleaning the area with antiseptics, a local anaesthetic (lidocaine) was injected on the skin over the skull, a longitudinal incision was made along the animal’s head and a small craniotomy was performed over the ACC (0.75 mm rostral and 0.4 mm lateral from Bregma). A glass micropipette containing viral reagents was positioned in place (1.75 mm depth from skull) and an air pressure system (Picospritzer II, Parker Hannifin, NH, US) was used to deliver ~500 nL of volume at a rate of ~100 nL / minute. After injections, glass micropipettes were live in place for another 5 minutes to allow the virus to diffuse at the injection site and then were slowly removed. Finally, the skin was sutured with an absorbable thread (4-0 coated VICRYL rapid suture, Ethicon). Additionally, a mix of analgesics and anti-inflammatory was subcutaneously injected (Dexamethasone, 2 mg/Kg; carprofen 5mg/kg). Animals were allowed to recover from the surgery and weight-checked every day for one week.

#### 1.2. GRIN lens implantation

Grin Needle Endomicroscope lenses of 1mm diameter, 4.38 mm long, 1:1 magnification and non-coated (NEM-100-25-10-860-S) were purchased from GRINtech (Jena, Germany). Seven to 14 days after virus injections, mice were anaesthetized as indicated above, placed in a stereotaxic frame and maintained their body temperature at 35°C. Eye ointment (Pharma Medica AG, Rogwil, Switzerland) was applied and antiseptics and local anaesthetic (lidocaine) were administered on the skin over the skull. After head hair removal and opening the mouse skin, the surface of the skull was cleaned and dried and a thin layer of Cyanoacrylate glue was applied to ensure a better adherence for cementing the lenses. Then, a circular craniotomy of just the size of the lens was drilled at the virus injection site over the ACC and remaining bones and dura carefully removed. Surgical area was maintained wet with artificial cerebrospinal solution until the end of lens implantation. Grin lenses were descended to its final position (~1.5 mm depth from skull) grabbed by a homemade holder attached to a micromanipulator. Lenses were lowered at a speed of 20 microns every 30 seconds. This procedure minimizes resistance to penetration into brain tissue and prevents increased intracranial pressure. Afterwards, the skull was dried with synthetic surgical sponges (Fine Science Tools) and a self-cure dental adhesive (Super-Bond C&B, Sun Medical, Japan) was applied to cement the lenses. After adhesive curated and lens holders were carefully removed, plastic cylinders made of eppendorf tubes were glued around the lenses to protect them from scratches. Finally, a metal bar serving to clamp the mice on a holder under the two-photon microscope was cemented (Paladur, Heraeus Kulzer GmbH, Germany) to the posterior part of the skull. Additionally, a mix of analgesics and anti-inflammatory was subcutaneously injected during the surgery (Dexamethasone, 2mg/Kg; Carprofen 5mg/kg). Animals were allowed to recover from the surgery and controlled on a regular basis. Moreover, animals were chronically injected for 7 days with a mix of Dexamethasone (0.5 mg/kg) and Carprofen (5mg/kg).

#### 1.3. Chronic constriction injury and Von Frey test

Sciatic nerve lesions and Von Frey tests were carried on as previously described [28, 29, 32]. Mice were anesthetized with isoflurane as indicated above and surgery was performed under a sterile hood. Eye ointment (Pharma Medica AG, Rogwil, Switzerland) was applied and fur was shaved from the left thigh. A small incision was performed with a scalpel and the sciatic nerve was reached by separating the two muscles above. The muscles were kept apart with a small tissue retractor (Fine Science Tools, Heidelberg, Germany, and a sterile sofsilk thread 5-0 (Coviden, Dublin, Irleand) was used to make 3 consecutive loose ligations (~1 mm apart) on the nerve. The skin was then sutured with an absorbable thread (4-0 coated VICRYL rapid suture, Ethicon). Animals were allowed to recover from the surgery and controlled on a regular basis. For Sham surgery an equivalent parallel operation was performed on littermates, in which the sciatic nerve was exposed but not ligated.

Mechanical threshold was evaluated with the Von Frey test one day before and 7 day after sciatic nerve surgery. Mice were placed on an elevated grid inside a Plexiglas cylinder and allowed to habituate for 20-30 minutes. Mechanical hyperalgesia was then tested using an Electronic von Frey aesthesiometer (IITC Life Science, CA, USA) by slowly applying pressure to the midplantar surface of the hind-paw with the Von Frey filament until a paw withdrawal was evoked. Six pressure measures were taken per each paw and an unpaired t-test was used as exclusion/inclusion criterion.

### 2. In vivo two-photon calcium imaging

Animals were imaged under light isoflurane (0.5-0-9%) anaesthesia using a custom-built two-photon laser-scanning microscope [33] with a 20x water immersion objective (W Plan-Apochromat 20x/1.0 DIC VIS-IR, Zeiss). GCaMP6 was excited at 910 nm with a Ti:sapphire laser (Mai Tai; Spectra-Physics), and emission was detected with GaAsP photomultiplier modules (Hamamatsu Photonics) fitted with 520/50 nm band pass filter or a 607/70 band pass filter and separated by a 560 nm dichroic mirror (BrightLine; Semrock). The microscope was controlled by a customized version of ScanImage (r3.8.1; Janelia Research Campus).

The first imaging session was performed 3 weeks after lens implantation. For this and further sessions, a digital zoom of 3 was used. Focusing on a border of the grin lens was used as a landmark for navigating in x, y and z axis. High resolution images (512 x 512 pixels, 1.11 Hz) were collected to identify an appropriate field of view (as many cells as possible). Then, a 3 minutes high resolution movie was saved as reference to identify the same cells in successive sessions. Spontaneous and stimulus-evoked calcium transients were recorded using fast images (256 x 128 pixels, 8.49 Hz). All sessions consisted of 10 minutes of spontaneous activity followed by assessments of neuronal responses to a set of different stimuli. Heat and warmth stimuli consisted of one minute trials repeated 10 times, whereas all other stimuli comprised 20 seconds trials repeated 30 times. All trials included 10 seconds of pre-stimulus neuronal activity for statistical comparison with stimulus-evoked responses. All stimuli (otherwise stated) were applied to the hindpaw ipsilateral to the operated sciatic nerve (contralateral to the site of imaging). Stimulus modalities tested were as follows:

Hindpaw electric shock (HPS): a short electrical pulse (50 ms; 90 V) delivered to the hindpaw. Electrical stimulation was achieved by applying a brief current onto conductive adhesive strips (approximately 1 cm wide by 2 cm long) placed on the hindpaw pad. Forepaw electric shock (FPS): same as HPS but applying the conductive adhesive strips on the forepaw (ipsilateral to the operated sciatic nerve). Heat: noxious heat (48°C) applied to the hindpaw. A custom-made heating device made of an aluminum plate attached to a peltier element was put in contact with the hindpaw pad. The aluminum plate was hold at 32° until a heating ramp was launched. The ramp consisted of a 16-seconds rise phase at a rate of 1°C per second, followed by a 6 seconds plateau at 48°C and a decay phase where the plate was cooled until it reaches 32°C again. Warmth: non-nociceptive warmth (43°C) applied to the hindpaw. Similar methodology used for heat stimuli. This time the warming ramp consisted of 11 seconds rise phase at a rate 1°C per second, 11 seconds plateau at 43°C, and a decay phase cooling the plate until it reaches 32°C again. Touch: tactile stimulus delivered to the hindpaw. A Von frey filament attached to a homemade piezo-driven mechanical stimulator was used to press the hindpaw pad. Pinprick: Same methodology as for touch, but the von Frey filament was changed by a 30G needle. Sound: loud noise as an aversive but non-nociceptive sensory stimulus. A white noise generator was used to deliver a loud tone of 72 dB (500 ms duration; background noise: 58 dB). Puff: aversive but non-nociceptive air puff (500 ms, 20 psi) delivered to the cheek-eye region through an air pressure system.

### 3. Calcium imaging data analysis

We processed all calcium imaging data using a MATLAB-based custom-made pipeline. However, certain routines were run on either Python or R as stated below.

#### 3.1. Image registration, motion correction, fluorescence extraction and spike inference

After acquiring all sessions for a given animal, calcium imaging videos were registered and motion corrected using the NoRMCorre algorithm in MATLAB software environment [34]. Once all frames were registered, we calculated the projections (mean, median, max, std, and cross-correlation) of the motion corrected movies which were later passed to our custom segmentation GUI written in Python. This implementation allowed us to select stack projection of frames of interest (from few frames to all frames present in concatenated sessions) and visually identify neurons through the cross-correlated pixels. We did not apply neuropil subtraction. We manually selected few regions of interest (ROIs) as landmarks to further perform minor adjustments to the field of view (FOV) from different sessions, namely rotation, stretch and squeeze, until all FOVs were aligned and passed quality-check. Once that was achieved, we continued with the segmentation of all ROI in the aligned FOV. After extracting the fluorescence of each ROI, we calculated ΔF/F using a running-window average of the 8th percentile of each ROI’s fluorescence using a Matlab toolbox, described previously [35].

For many of the analysis described below, we used the most likely discretized spike train underlying a fluorescence trace. This detection of calcium event-related spikes for each ROI consisted in a three-step method. First, spikes were inferred using the OASIS toolbox in Matlab [36], where we used the *‘constrained foopsi’* method and time constants were automatically detected.

Second, calcium events were automatically detected using continuous wavelet transform (CWT) implemented in R. Third, we then compared the events obtained by the CWT and the spikes by OASIS and we considered only spikes that did fall under a detected calcium transient. Denoised traces were computed from the convolution of the spikes using rise-time kinetics of 1 ms and decay time kinetics of 380ms.

To assess the spontaneous activity of a given ROI, we computed the mean event amplitude and mean event rate from the ΔF/F traces for each ROI in a total of 10 min recording (see above). Single events were detected using the method described above. We computed the peak amplitude by taking the difference of max ΔF/F and the baseline. We averaged the peak amplitude and the event rate per ROI and we pool together all ROIs for all recorded animals.

#### 3.2. Analysis of stimulus-evoked responses

In order to detect responsive neurons, we hypothesized that the neuronal activity after stimulus must be significantly different that of baseline. To test this hypothesis, we used logistic regression as described below. We used the *nxfxt* matrix from the spike inference protocol (see above), where *n* = number of cells, _*f*_ = number of frames per trial, and *t* = number of trials. Then, in order to speed up computations, we binned the data by averaging 4 consecutive frames (0.5 seconds). The baseline and the evoked window consisted in 11 bins (5 seconds) each. Given the nature of the temperature stimuli (heat and warmth) the time at which we consider the evoked window differs from the rest of stimuli. In the case of heat, we considered 16 seconds after the onset of the peltier element and for warmth we consider 11 seconds after the onset of the peltier element. In both cases the evoked window was the same that of the rest of stimuli (5 seconds). After binning the spikes, we vectorized them per ROI and classified the responses using logistic regression by the Scikit-learn machine learning library in Python. We thought that if the neuronal activity is different in the evoked window, it will result from significantly modulated bins. To detect bins that showed modulatory activity, we first assessed the difference from chance. Briefly, we performed *k*-fold cross-validation, with *k* = 5, so that in each iteration 80% and 20% of the data were used for training and testing respectively. We obtained posterior probabilities of the observations per bins through all trials and we compared it to a random (null) distribution. We assessed statistical significance by bootstrapping the data using 1000 permutations, obtaining the p-value (p_boostrap) of that difference. We considered only bins that showed a significant difference of p_boostrap < 0.01. If we detected bins that differ from chance, we next compared the activity of those bins of activation to the baseline period. We first computed a confusion matrix based on true positives (*TP*), false positives (*FP*) and false negatives (*FN* = *n*_(*pos*)_ – *TP*, where *n*_(*pos*)_ describes the number of positive observations). Using this confusion matrix we calculated the Precision (*P*) & Recall (*R*) values as follows:

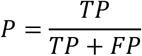

and,

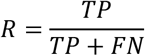

and then we calculated the area under the Precision/Recall curve (*AUPRC*), and we normalized it to an interval between 0 and 1, where chance = 0.5, using the following formula:

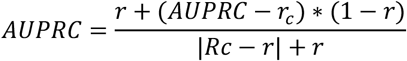

where r denotes chance at 0.5, *r_c_* is a random classifier given by the umber of observations as,

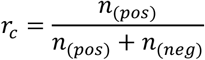

where *n*_(*neg*)_ denotes the number of negative observations

We then used permutation test to compare the values of AURPC to a null distribution using shuffled baseline and evoked activity values, We considered significance if the p-value (p_AUPRC) < 0.01. We considered that a neuron showed significant modulation only if p_boostrap < 0.01 and p_AUPRC < 0.01. We classified a neuron as being activated by the stimulus when AUPRC > 0.5, and a neuron being inhibited when AUPRC < 0.5. We named these values as ‘selectivity’. Therefore, the term ‘selective’ denotes a neuron that has significant modulation with values of AUPRC different than 0.5. We assessed the *response probability (reliability*) by the sum of significant trials divided by the total number of trials for the given stimulus. We calculated the *proportion of responding cells* to a given stimulus by taking the number of neurons that were activated or inhibited over the total cells imaged. We also provide the estimate of proportion of neurons being modulated by one or more stimuli. For it, we count the number of neurons that showed significant modulation to different stimuli over the total number of cells.

##### Identification of nociception, aversion and saliency-specific neurons

for each session, all neuron were classified according to its responsiveness to the set of stimuli tested as: i) non responsive; ii) discrete: responsive to only one out of all stimuli; iii) nociception-specific: activated exclusively by two or more noxious stimuli; iv) aversion-specific: responding exclusively to two or more aversive (noxious or not) stimuli; v) saliency-specific: activated by two or more stimuli. Categories iii, iv and v are accumulative, as a nociception-specific cells are also aversion and saliency-specific while aversion-specific are also saliency-specific. The proportions obtained for each category were compared to a chance distribution. For this, we generated 1000 of simulated response matrices by shuffling, for each stimulus, the identity of the responsive and non-responsive cells keeping constant the proportion of neurons being activated. Then, for each category we calculated a *z-score* as follows:

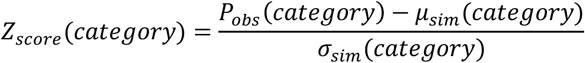

Where for P_obs_ are the proportion of cells observed in the real data and μ_sim_ and σ_sim_ are the mean and standard deviation of the proportion of cells obtained in simulated response-matrices. Statistical significance was evaluated from *z-score*s with a threshold of p= 0.05.

To assess the stability of this representation, we identified cells conserving category-specificity from one session to the next and compared the obtained distribution to simulated ones. For this, we generated 1000 simulated response matrices based on real data from session 2 and compared each of it with the real response matrix from session 1. Then, for each category, we calculated a *z-score* using the observed and simulated proportion of neurons conserving specificity across sessions.

##### Overlap representation

The longitudinal analysis performed to evaluate the overlap of touch and HPS, and touch and heat was performed at 14d after surgery.

##### Response amplitude

We calculated the mean (*μ*(*bsl*)) and standard deviation (*σ*(*bsl*)) neuronal activity for 5 s before the stimulus onset and computed the z-*score* for the 5 s before and after the stimulus using the following formula:

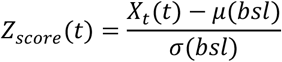

Where *t* denotes the frame relative to the stimulus onset, and *X* denotes the ΔF/F of an individual trial.

##### Time-lapse stimulus representation

In order to evaluate whether neurons are activated by the same stimulus over sessions, we selected the activations by heat, hind-paw electric shock and air puff. For each timepoint, we calculated the change in representation per animal (D) as follows:

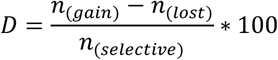

where *n*_(*gain*)_ denotes the total number of cells that gained selectivity, per timepoint (not responding in first session and responding in second session), *n*_(*lost*)_ indicates the number of cells that lost selectivity per timepoint (responding in first session and not responding in second session), and *n*_(*selective*)_ denotes the number of cells selective in either session per timepoint (i.e. responding in either the first session OR the second session).

We then grouped all the animals and split them into their respective experimental group and we tested the hypothesis that, even though there is a change in neuronal responsiveness to a stimulus, the global representation does not change. If D is positive, it indicates that there are more cells that gained selectivity than lost selectivity; and vice versa. For testing this hypothesis we computed one-sample t-test. We also performed statistics on cross-sectional D values. For each time point we tested the distribution of data points of sham animals and CCI animals. We conducted un-paired two-samples t-test.

#### 3.3. Stimulus decoding

To evaluate the neuronal code of stimulus response, we implemented Logistic Regression classifier using the Scikit-learn machine learning library in Python as follows: For each recording session, we binned the *n* x *f* x *t* matrix of the evoked period in 25 frames (3 seconds) each, and we averaged the spikes found in each bin, obtaining a *n* x *b* matrix, where *n* is the number of cells and *b* is the binned data. We assumed independence between each bin. As stated above, the time window we took for warmth and heat are 11 and 16 seconds after the onset of the Peltier element. The matrix then was vectorized in 1 x *b*. We assigned each stimulus a class label and we split the data into testing and training sets per trial with 0.8 and 0.2 randomly selected fractions, respectively, 5 times. We performed k-fold cross validation. Each time, we trained the classifier with a subset of the bins, and we estimated the posterior probabilities for each bin belonging to each class. Using the actual labels and these respective probabilities, we then computed the Precision / Recall curve while comparing the data to two distributions (as described below). Briefly, first we computed a confusion matrix of the tested data subset using the PyCM modules in Python. The confusion matrix indicates the number of correct classifications of each bin into the respective label given the probabilities calculated above, where rows indicate predicted labels and columns indicate actual labels. From these matrices we obtained the values for AUPRC for each class. At the same time, we collected the confusion matrices for further analysis. The values of the AUPRC were normalized to 0-1 interval, where 0.5 indicates chance, as stated above. We then normalized the confusion matrix values by the number of classes for each train/test split (total of 5 splits) to obtain a matrix with values from 0 to 1 where the sum of predicted values adds up to 1. After all the splits were completed, the values of the confusion matrices as well as the values of the AUPRC were averaged. We assessed the performance of the classification for the actual stimuli by estimating the predicted stimuli for each confusion matrix by comparing the correct classification percentage to that of the incorrect classified stimuli (i.e. off-diagonal). We tested for significance using Wilcoxon sign-rank. We then hypothesized that chronic pain might lead to an increase in the number of incorrect classifications of an innocuous stimulus (Figure 4C). To test this hypothesis, we grouped stimuli according to their nature, where air puff and sound were considered as aversive stimuli, pinprick, heat, HPS and FPS as painful stimuli (Figure 1F). Using this categorisation, we compared the values of the error predicted classification (false-positives) for an actual stimulus or stimuli combination in chronic pain animals that of the sham animals using Mann-Whitney test. Additionally, we tested stability of the population code over time (generalizability). We calculated the posterior probabilities (as done above) of the data of day 2, using the trained classifier of day 1. We computed the confusion matrices using the PyCM library in Python as stated above using the data from day2. We computed the decoding accuracy by

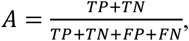

where *TP* = True positive, *FP* = False positive, *TN* = True negative, and *FN* = False negative.

#### 3.4. Clustering

We performed cluster analysis to evaluate the effect of chronic pain at two weeks after surgery on the quantitative population stimulus representation in a selectivity space. We hypothesized that chronic pain does not only alter the proportion of neurons responding to innocuous stimuli, but also it may alter how the neurons respond to the stimuli. This might be reflected in the selectivity value. Therefore, we aimed to characterized the relative distance of stimuli selectivity, performing dimensionality reduction and cluster analysis. We performed two separate sets of analysis.

First, we aimed to determine the relative distance between single stimuli in a two-dimensional space using the recently developed UMAP (uniform manifold approximation and projection for dimension reduction) algorithm [37]. The data consisted in a *n* dimensional vectors of pair of stimuli selectivity, where n is the number of cells from those animals where the given pair of stimuli were considered (612 cells in sham, 964 cells in CCI : HPS-Heat; 371 sham, 544 CCI : touch-HPS (same for touch-heat)). We analysed the following pairs: HPS - touch, heat - touch and HPS - heat. Second, we performed unsupervised k-means clustering on individual stimulus selectivity, with *k* = 2. We found the most significant clusters defined by the Silhouette and Elbow methods. In this case we performed k-means directly of the *n* x *m* (n = number of cells, m = number of stimuli present in that session) matrix per animal, per session, and calculated the Euclidean distance between the centroids for each cluster. We obtained the centroids of each cluster by choosing the most representative cluster for each stimulus, as a ‘win-takes-all’ constrain. After obtaining all the centroids and the pair-wise Euclidean distances, we categorized the stimuli into painful, aversive (non-painful), and innocuous. We then compared the difference between the distance of centroids for given categories between sham and neuropathic animals.

#### 3.5. Statistical analysis

We performed statistical analysis within the *Matlab*(2018b and 2020a), *Python3*, and *GraphPad Prism 8* environments. All data were tested for significance and corresponding parametric or no parametric tests were used. Significance, p < 0.05.

## Notes

### Competing Interest Statement

The authors have declared no competing interest.

